# Microbial Community Interactions on a Chip

**DOI:** 10.1101/2022.10.17.511674

**Authors:** Duane. S. Juang, Wren E. Wightman, Gabriel L. Lozano, Layla J. Barkal, Jiaquan Yu, Manuel F. Garavito, Amanda Hurley, Ophelia S. Venturelli, Jo Handelsman, David J. Beebe

## Abstract

Multispecies microbial communities drive most ecosystems on Earth. Chemical and biological interactions within these communities can affect survival of individual members and the entire community. However, the prohibitively high number of possible interactions within a microbial community has made the characterization of factors that influence community development challenging. Here we report a Microbial Community Interaction (μCI) device to advance the systematic study of chemical and biological interactions within a microbial community. The μCI creates a combinatorial landscape made up of an array of triangular wells interconnected with circular wells, which each contains either a different chemical or microbial strain, generating chemical gradients and revealing biological interactions. *Bacillus cereus* UW85 containing GFP provided the “target” readout in the triangular wells, and antibiotics or microorganisms in adjacent circular wells are designated the “variables”. The μCI device revealed that gentamicin and vancomycin are antagonistic to each other in inhibiting the target *B. cereus* UW85, displaying weaker inhibitory activity when used in combination than alone. We identified three-member communities constructed with isolates from the plant rhizosphere that increased or decreased growth of *B. cereus*. The μCI device enables both strain-level and community-level insight. The scalable geometric design of the μCI device enables experiments with high combinatorial efficiency, thereby providing a simple, scalable platform for systematic interrogation of three-factor interactions that influence microorganisms in solitary or community life.

## Introduction

Much of modern-day microbiology was established from the one microbe-one disease hypothesis known as Koch’s postulates, formulated by Robert Koch and Friedrich Loeffler in the late 1800s, which generated 150 years of research on microorganisms living in solitary culture. Therefore, most knowledge of microbial behavior is derived from organisms living in isolation. Despite the enormous advances facilitated by pure-culture studies, they ignore the highly social nature of most microorganisms, which reside within complex multi-species communities. These communities play significant roles in human and environmental health. Although the classic approach of studying individual microbes in isolation offers advantages of simplicity, it cannot recapitulate the phenotypes that arise from microbial community interactions. Microbes within a community can exchange nutrients, metabolites, and information with each other via signaling with diffusible factors (e.g., quorum sensing), and these interactions drive the structure, dynamics, and phenotypes of microbial communities.

Microbial communities determine the health of their hosts. Soil communities are critical for the health and productivity of plants (1–4) and in the prevention of plant disease. For instance, disease suppressive soils (5, 6), defined as soils that suppress the establishment or persistence of soilborne pathogens, are commonly established via the enrichment of certain microbial species which act synergistically as a part of a beneficial microbial consortium against the pathogen (7–9). The discovery of this natural plant defense mechanism has evoked interest in developing microbial communities as biological control agents (10, 11). In human health, there is growing attention to the role of microbial communities in disease, which has spurred efforts to engineer microbial communities for therapeutic interventions (12–14). For instance, infection by *Clostridium difficile* (*C. diff*.) commonly emerges after antibiotic treatment, which reduces microbial diversity in the gut, opening up opportunities for pathogenic microbes such as *C. diff* to colonize (15, 16). Clinical trials also found that fecal microbiota transplantation was highly effective in treating *C. diff* infections (17), which have generated broad interest in developing more refined treatments. To design these interventions systematically, we need a better understanding of microbial behavior in communities. However, few model systems provide the means to dissect communities in a manner that parses variables to generate causal associations. Standard technologies for studying microbial communities include 16S ribosomal RNA gene (rRNA) sequencing (18, 19), fluorescence in situ hybridization (FISH) (20, 21), and macroscale (mostly pairwise) co-cultures (20, 21). 16S rRNA gene sequencing can provide valuable information regarding the composition of a community but cannot elucidate specific interactions among members. FISH can provide information regarding the spatial distribution of different microbial members within a sample but cannot provide functional readouts and is challenging to perform. Pairwise co-cultures are generally performed in conventional laboratory culture vessels like multi-well plates, tubes, and solid agar plates. However, they are generally not well-suited for systematically screening interactions that occur between more than two microbes, and miss emergent properties present between three or more members within the community (22–27). A significant bottleneck in microbial community studies is the lack of simple, practical, and scalable co-culture tools to enable systematic functional screening of microbial community interactions. This capability is essential to enable scientists to gain functional insight on how different community members interact and drive community behavior within the microenvironment, which is fundamental both in the study of natural microbial communities and in rationally designing synthetic microbial communities.

More recently, there have been developments in employing microfluidic technologies for the study and assembly of microbial communities. The majority of these studies employ droplet microfluidic technology, i.e., using fluorescently encoded droplets (or microbes) for performing random community assembly by droplet merging (28–31). Microbial growth is monitored using fluorescent reporter strains or fluorescent metabolism tracking dyes. Droplet microfluidics offers unparalleled throughput (>10^6^ combinatorial conditions), but the technology still has a few limitations, including (1) difficulty in quantifying the growth of strains in a mixed-species community in a given droplet unless different fluorescent reporter strains are used, and (2) limited culture time due to the small volume (∼nL scale) of droplets. Although a few non-droplet-based microscale co-culture platforms have also been developed for microbes (32–36), these approaches are seldom designed to examine the interactions of more than three microbial members at one time and are often relatively complex to operate, requiring sophisticated microfluidic designs or 3D bioprinting techniques.

Here we report a simple and scalable microbial community co-culture platform termed the **<u>M</u>**icrobial **<u>C</u>**ommunity **I**nteraction (μCI) device. The μCI device enables spatially segregated co-culture of several interacting microbial members at the same time, allowing monitoring the growth of each member independently, but limits the interrogation of interactions to a “localized” limited parameter space for ease in data interpretation. This provides a community-level approach to functional studies but preserves strain-level resolution for interactions. We first demonstrated the combinatorial screening capability of the μCI device via the screening of antibiotic combinations and identified an antibiotic combination (gentamicin and vancomycin) which showed interfering activity in inhibiting the growth of *Bacillus cereus* UW85 (*B. cereus* UW85). We further developed a custom software pipeline to automatically generate an optimal combinatorial experimental design layout and automated image data analysis. Lastly, using *B. cereus* and ten rhizosphere microbial co-isolates from the soybean rhizosphere as a model, we demonstrated the utility of the μCI device for screening community-induced changes in the survival of *B. cereus* as a target microbe.

## Results

### Design of the μCI device

The μCI device was developed with the following design principles (1) it must be compatible with current standard microbial culture methods (agar plates), reagents (standard microbial culture media such as LB), and readouts (fluorescence microscopy/visual inspection) and allow for the control over culture conditions; (2) simple to operate and implement by the microbiological community using only readily available standard laboratory equipment; (3) possess a simple and high content data readout; (4) easily scalable in terms of the number of microbial members. The core design of the μCI device is an array of equilateral triangular combination wells, which are each connected to three circular variable wells by each vertex of the triangle. Each circular well can contain a different variable, and each is connected to up to six triangular combination wells in a hexagonal layout (Fig. 1). The variable wells can each contain a different experimental treatment, such as a microbe, antibiotic, or other factor, whereas each triangular combination well can contain a fixed factor, a microbial “target” strain. In this way, each triangular combination well can be considered a function of the sum of three variable wells. This design enables the systematic screening of three interacting variables within one well simultaneously.

**Fig. 1.**
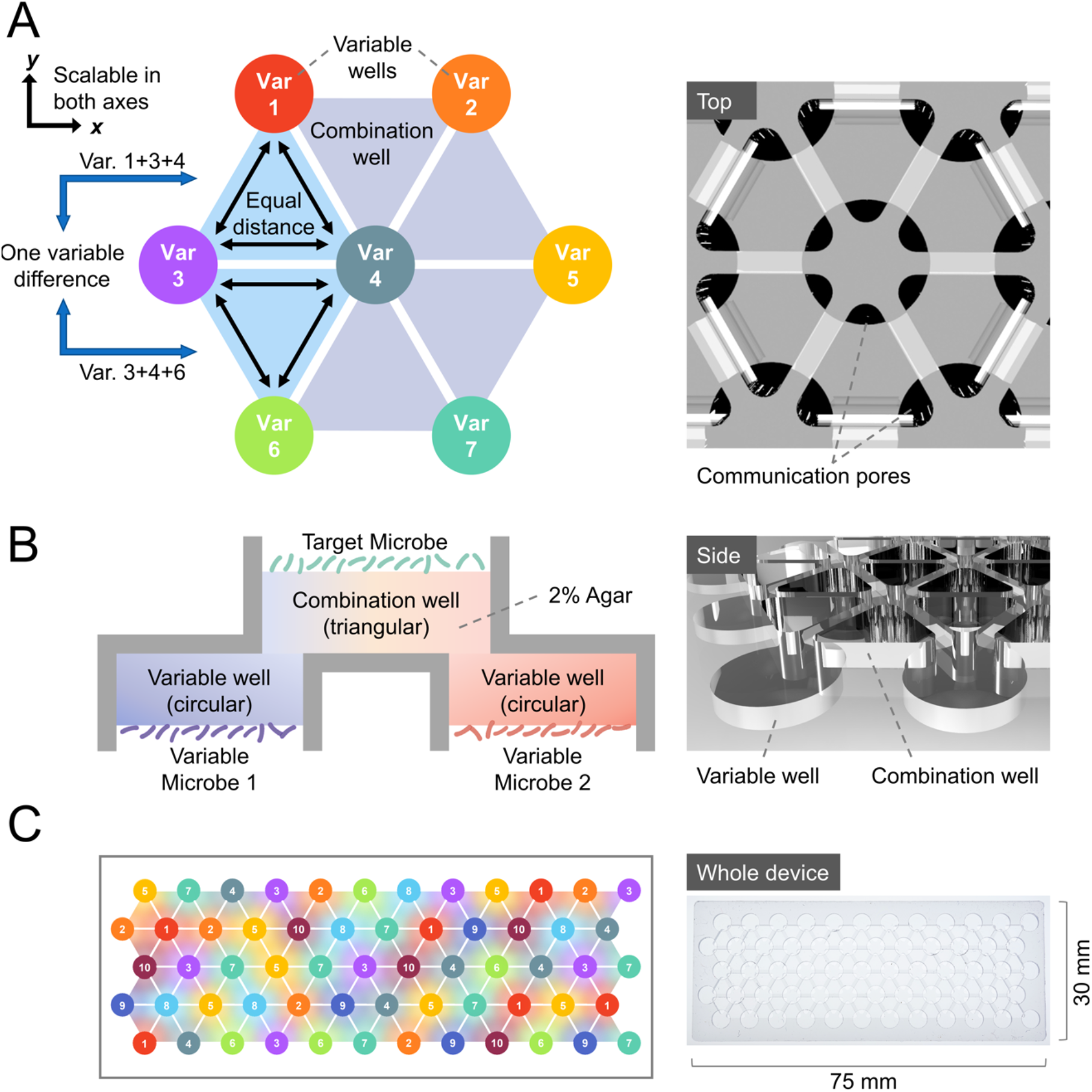
Design of the μCI device. (A) Top view, (B) Cross section, and (C) Whole-device schematics and images of the μCI device. Each combination well generates a three-variable combination gradient.

The choice of the triangle instead of other geometrical configurations is due to the following considerations (1) The distance between each vertex of an equilateral triangle is the same, which allows for “equal strength” in terms of interaction distance between different microbial members, which cannot be accomplished using geometries with larger number of sides; (2) The triangle is the most basic unit that can define a planar surface, which allows the community interaction landscape to scale out in both the *x* and *y* dimensions, whereas a linear two-way interaction network is confined to only one dimension at a time in terms of geometrical design; (3) A triangle can be joined with another equal triangle via one side, which allows the two triangles to share two connecting vertices, but each having one vertex that is not shared. This allows for a three-factorial but single variable comparison between any two neighboring triangular combination wells; and (4) three-way interactions are more manageable in terms of scale of experiments compared to higher numbers of combinations for the number of participants within a microbial community.

The μCI device is fabricated out of polystyrene and consists of circular variable wells on one side and triangular combination wells on the other side, facing opposite directions (Fig. 1). The overlapping opening area (communication pores) between the variable and combination wells allows contact-independent communication through agar gel (Fig. 1A). Although microbial interactions can be contact-mediated, it is challenging to achieve spatial isolation (thus enabling individual optical readouts of growth for each member of a community) using contact-mediated mixed co-cultures. To optimize the design of the μCI device for testing microbial communities, we tested various device sizes with a well pitch (center to center distance in the *xy* plane of the nearest neighboring well) ranging from 4.5 mm to 6.5 mm, but with the same thickness (*z*) across all devices, to find an optimal diffusion distance for co-culture, as microbial interactions within the μCI device are diffusion mediated. That is, a diffusion distance that is short enough for the target strain to show robust growth in a community co-culture within a 1∼2-day culture period, but not too short so that cross-combination well interference may become an issue. Each variable and combination well holds a given volume of solid agar, so secreted factors are also diluted when diffusing across the agar, with larger wells (longer well pitch) diluting the secreted factors more. We first characterized soluble factor diffusion through the device using a fluorescent small molecule with a molecular weight similar to that of antibiotics (Rhodamine 6G, MW: 479.02) with time-lapse fluorescence microscopy to monitor the diffusion profile over time (Fig. S1A, S1B). Results show that the fluorescent small molecule was more rapidly depleted from the 4.5-mm pitch device, and more slowly from the 5.5-mm, 6-mm, and 6.5-mm device (Fig. S1C, S1D) as is expected due to its shorter diffusion distance. To further investigate the diffusion behavior of the μCI device, we added a high concentration antibiotic (80 μg/mL gentamicin, MW: 477.6) to one of the variable wells, and inoculated *B. cereus* UW85 containing a plasmid that carries the gene encoding green fluorescent protein (GFP) (*B. cereus* UW85 GFP) as a target strain to the combination wells (Fig. S2A). Fluorescent images of the combination wells were captured following 1-day of incubation. Results suggest significant cross-combination well interference in the 4.5-mm pitch device (significant growth inhibition for neighboring combination wells), slight inhibition at the very edge of a neighboring combination well in the 5.5 mm pitch device, and no observed neighboring well inhibition for the 6-mm and 6.5-mm pitch devices (Fig. S2B). As such, we selected the 6-mm pitch design for all subsequent experiments. We also loaded four fluorescent cell-tracking dyes into each variable well of the 6-mm device to show that the device can successfully generate a three-factor combination gradient within each combination well (Fig. S1E). Finite element modeling (FEM) using COMSOL Multiphysics software also suggested that a gradient can be maintained across the agar surface of the combination well of the 6-mm device (Fig. S3A, S3B), and the FEM diffusion profiles appear similar to the fluorescence diffusion profiles of Rhodamine 6G (Fig. S3C). After optimizing the design of the basic hexagonal μCI unit, we scaled the hexagonal unit to a device with 60 variable wells and 88 combination wells (Fig. 1C) and another with 19 variable wells and 24 combination wells (Fig. 3, Fig. S4). It’s worth noting that different device configurations can be easily made by scaling the modular hexagonal unit to fit the scale of the experiment (number of combinations required).

**Fig. 3.**
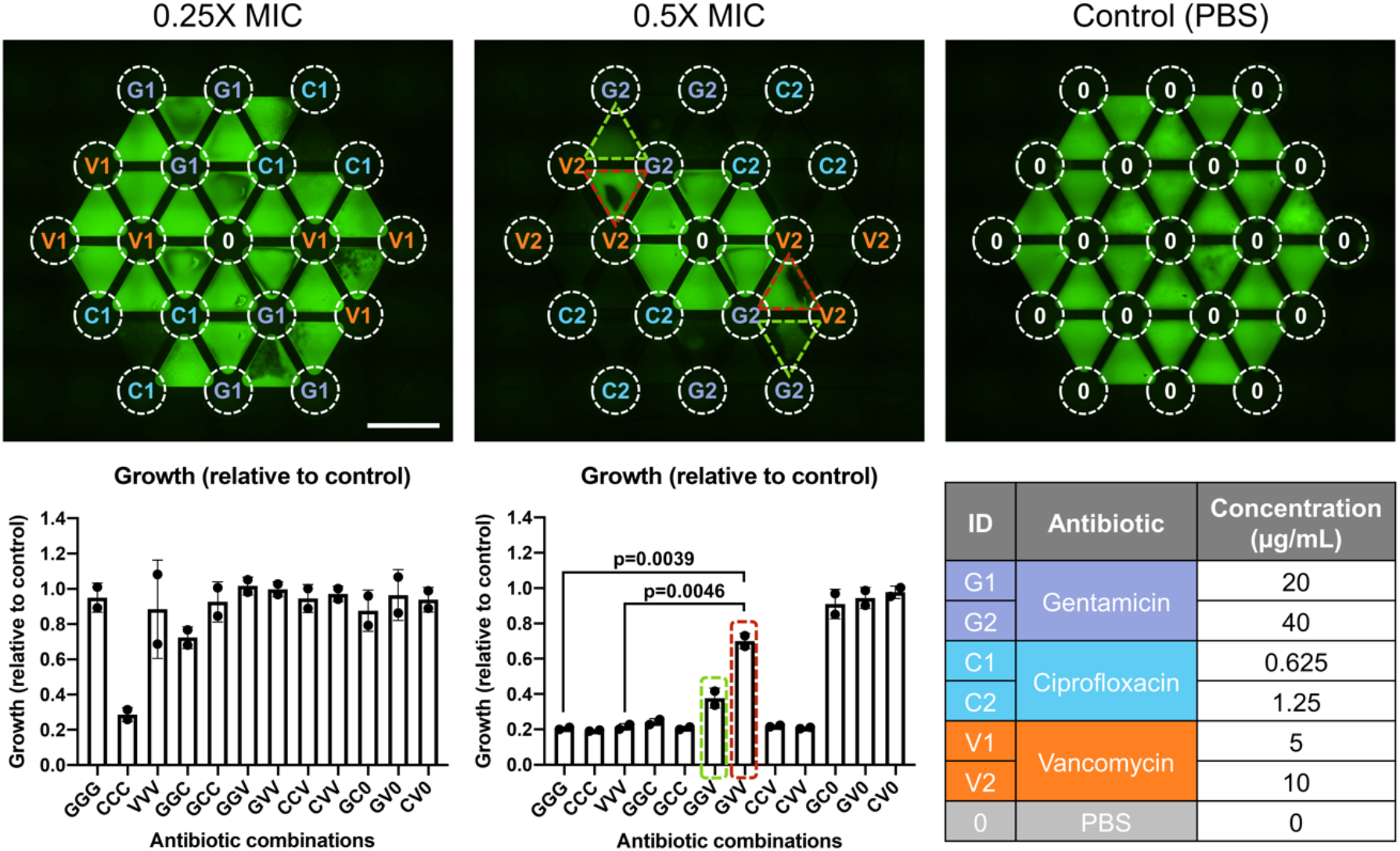
Antibiotic combination testing on a 24-combination well μCI device. Upper panel shows the fluorescence microscopy images of target strain *B. cereus* UW85 GFP on the μCI device after culture in the presence of antibiotics at various combinations or without antibiotics (control). Scale bar: 5 mm. Red and green dotted lines highlight the wells displaying antagonistic antibiotic activity. Quantified growth (relative to no-antibiotic control) of the target strain when cultured in the presence of antibiotics at various combinations is shown in the lower panel. Red and green boxes highlight the antibiotic combinations displaying antagonistic activity (quantified from upper panel). Error bars denote the standard deviation from two replicates. Lower right legend shows the list of antibiotics and their concentrations used in this experiment.

### Experimental design and workflow of the μCI device

To manage the complexity of the set-up and data generated by three-factorial combinatorial experiments for large numbers of microbial strains, we developed a custom automated experimental design and data processing workflow in MATLAB (The MathWorks, Natick, MA) for the μCI device (Fig. 2). In brief, a user inputs the number of variables (microbial strains, antibiotics, etc.) to be tested for a given experiment into the MATLAB script, which attempts to search for an experimental layout with maximal combinatorial space coverage on the device and minimal repeats (details in Materials and Methods section). Microorganisms are then inoculated in the variable wells according to the generated layout, followed by inoculation of fluorescent target strains in the combination wells. After inoculation, fluorescence images of the target strains in the combination wells (facing down) are captured at set time intervals. The images are analyzed using a custom ROI array in Fiji (https://fiji.sc/) to quantify the mean fluorescence intensity of each combination well (which informs the total growth of the well) and the three vertices of the combination well (which informs which variable likely contributed more to the increase or decrease in growth of the target strain). The analyzed fluorescence intensities are then imported back into MATLAB, which maps each datapoint to the associated strain combination indicated by the generated experimental layout.

**Fig. 2.**
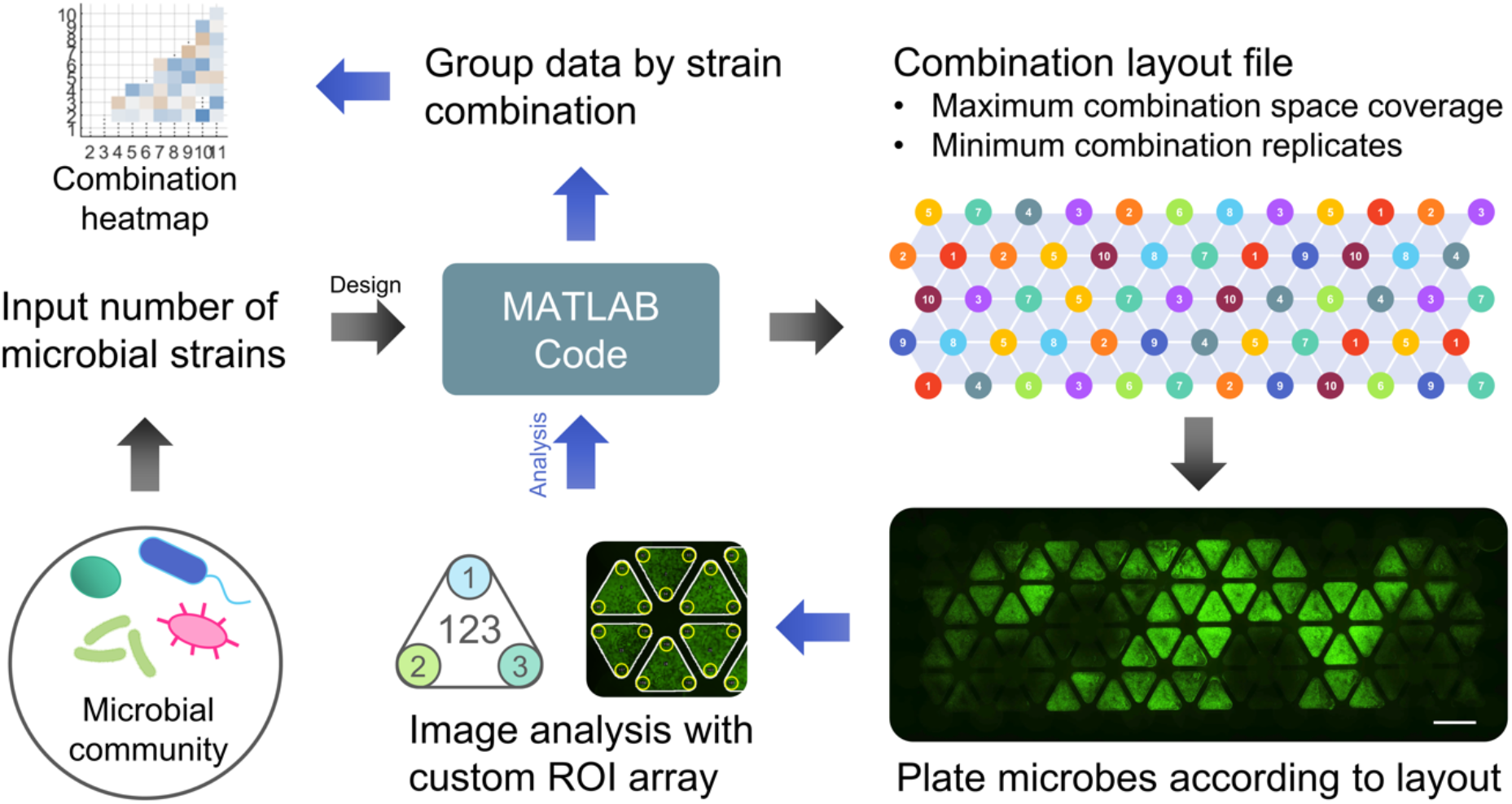
Experimental workflow of the μCI device. A custom MATLAB script generates random experimental layouts according to the number of microbial members to be tested, and for sorting and analyzing the quantified image data following experiments. ROI: region of interest. Scale bar: 5 mm.

### Antibiotic combination screening on the μCI device

To test the efficacy of the μCI device for multi-factorial experiments with small molecules, we selected three antibiotics: gentamicin, ciprofloxacin, and vancomycin, which inhibit growth of *B. cereus* (37–40). We first determined the minimum inhibitory concentration (MIC) of the antibiotics in the μCI device. Two-fold serial dilutions of the antibiotics were added to *one* of the three circular variable wells connected to a combination well on the μCI device. The reason for only adding antibiotics into one instead of all three variable wells is so that the gradient of antibiotic inhibition inside the combination well can be better observed. An overnight liquid culture of the target *B. cereus* UW85 GFP was diluted to OD 0.01 and inoculated into the combination wells. The devices were incubated at RT for two days with fluorescence images captured on both day 1 and day 2 to quantify the growth of *B. cereus* UW85 GFP. We defined the MIC as the antibiotic concentration in which a zone of growth inhibition was observed in the triangular combination wells adjoining the antibiotic-containing variable well (Fig. S4). MICs were 80, 2.5, and 20 μg/mL for gentamicin, ciprofloxacin, and vancomycin, respectively (Fig. S4). Worth noting is that this is the concentration of the antibiotic solution added to *one* of the three variable wells adjoining a combination well, not the actual antibiotic concentration the microbes are exposed to, which is hard to estimate due to the diffusion gradient nature of molecules in the μCI device. We then performed a three-antibiotic combination screening experiment on the μCI device at 0.25X and 0.5X the MIC of each antibiotic alone (Fig. 3). A control device (variable wells added with vehicle control (PBS)) was also run in parallel for growth normalization. Worth noting is that for the single antibiotic MIC experiment (Fig. S4), only one of the three variable wells adjoining a combination well contained antibiotics to facilitate the observation of an inhibition gradient, whereas in the three-antibiotic combination experiment, antibiotics were added to all three variable wells (or two of the three). Interestingly, we observed that at 0.5X MIC, a combination of gentamicin and vancomycin was less inhibitory than the individual antibiotics alone (Fig. 3, center panel) and this effect was more pronounced with a gentamicin to vancomycin ratio of 1:2 (GVV) than 2:1 (GGV). These results suggest that gentamicin and vancomycin are antagonistic to each other in inhibiting the growth of *B. cereus* UW85, and highlights the ability of the μCI device in discovering emergent phenotypes from multi-factorial combination experiments.

### Inoculation time and population density affects response of the target strain

The outcome of microbe-microbe interactions can be affected by the concentration and sequence of inoculation (41–43). For instance, microbes show higher tolerance to antibiotics when they have already grown to a high density compared to when their densities are low (44). In the μCI device, there is a time lag between inoculation and when the microbes start “sensing” each other in the device as it takes time for secreted factors to diffuse from the variable strains through the agar to reach the target strains in the combination wells. Initially, we inoculated the variable and target strains simultaneously in the device, but did not observe large differences in target strain growth when they were co-cultured with different communities. This led us to hypothesize that the target strains may have already established significant numbers before they start sensing and responding to the secreted factors from the variable strains, and hence co-culture-induced changes in target strain growth were muted.

To identify culture conditions which can allow us to observe stronger co-culture-induced phenotypes, we selected four strains from a previously reported collection of rhizosphere bacteria (45). *B. cereus* UW85 from this bacteria collection was selected as the target strain because it is a member of a widely distributed species in soil and on roots (46), and has many ecological roles in the rhizosphere (47–49). Moreover, *B. cereus* strains commonly carry microbial “hitchhikers”– bacteria that co-isolate with *B. cereus* but only become apparent in culture after incubating seemingly pure cultures at 4 ºC for several weeks (50). The hitchhiking behavior of *B. cereus* co-isolates suggest that these microbes live in close proximity in natural ecosystems, lending biological relevance for studies of their interactions *in vitro*. Indeed, *B. cereus* and two co-isolates have become a highly informative model designed THOR (abbreviation of “**t**he **h**itchhikers **o**f the **r**hizosphere”) for dissecting community behavior (45).

We then performed a study on inoculation time and concentration using four strains with variable inhibitory activities against the same members: *Chryseobacterium* sp. CI02, *Flavobacterium johnsoniae* CI04, *Pseudomonas* sp. CI14, and *Achromobacter* sp. CI16 and a target microbe, *B. cereus* UW85 GFP. This generates a total of three three-variable combinations. We found much greater differences in target strain growth when the variable strains were inoculated 6 hours prior to the target strains compared to when they were inoculated simultaneously (Fig. 4). For the 24-hour time point, the target strain showed the greatest differences in growth between different communities when the variable strains were inoculated at 10 times the concentration, 6 hours prior to the target strain (variable strain OD = 0.1, target strain OD = 0.01), (Fig. 4B). For the 48-hour time point, the greatest differential growth was observed when variable and target strains were both inoculated at OD = 0.01 (Fig. 4B). Additionally, at the 48-hour time point, we observed growth greater than the no-variable control when the variable strains were inoculated at OD = 0.01 and 0.001, but all community combinations exhibit growth lower than the control when the variable strains were inoculated at OD = 0.1 (Fig. 4B right panel, purple bar). Notably, the inhibitory activity of CI14 against UW85 was previously observed using the classical spread-patch method (45) and was reliably replicated in the μCI device. Additionally, we identified and quantified positive growth interactions (such CI02, CI04, CI16 inoculated at OD = 0.01) that would be challenging to observe using classical methods. To maximize differences in the observed growth of the target strain, we inoculated the variable strains 6 hours prior to the target strains with an inoculation OD of 0.01 for both variable and target strains for subsequent experiments.

**Fig. 4.**
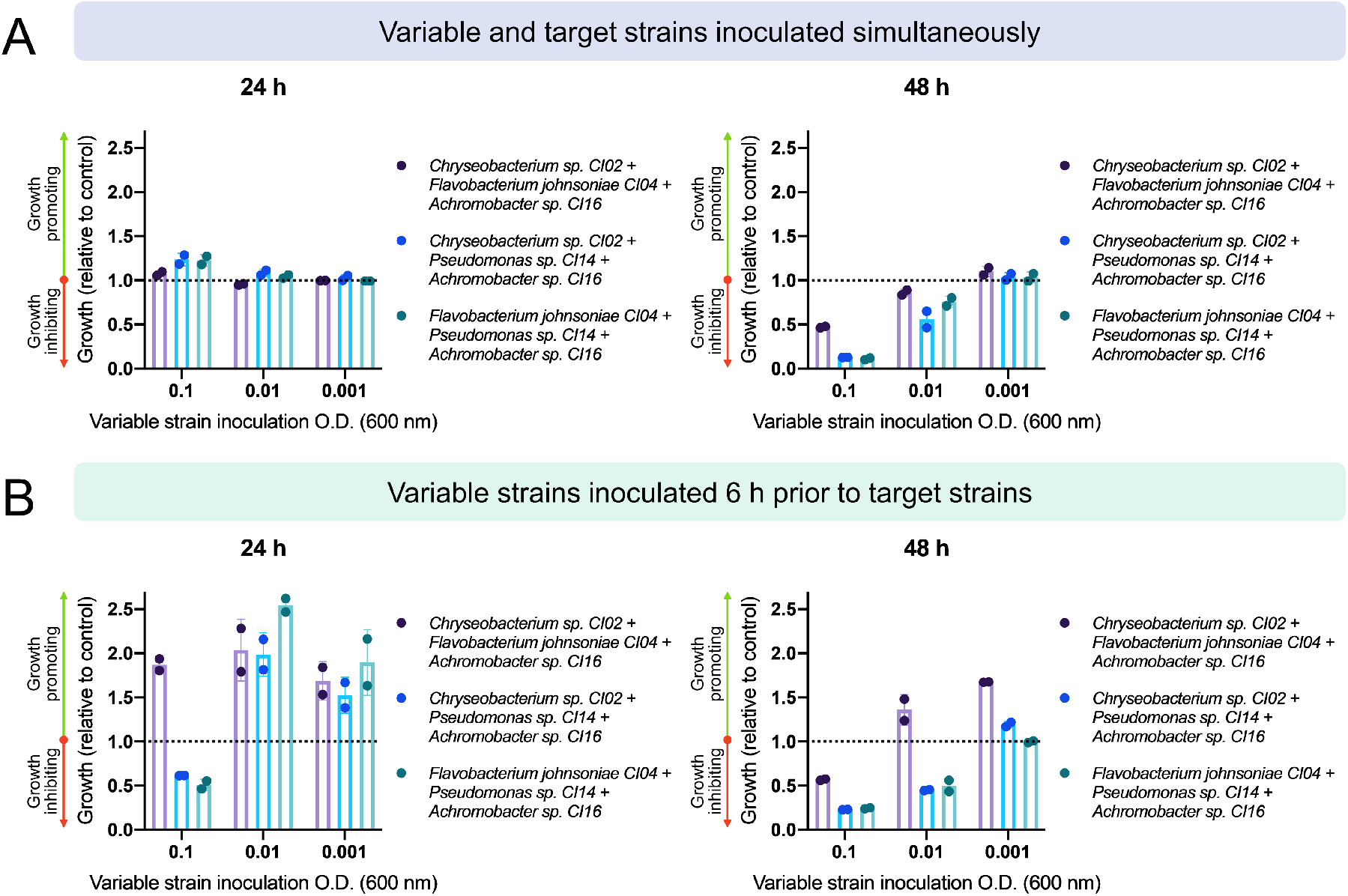
Inoculation time and concentration of variable strains affects growth response of target strain in the μCI device. (A) Variable strains (*Chryseobacterium sp. CI02, Flavobacterium johnsoniae* CI04, *Pseudomonas* sp. CI14, *Achromobacter* sp. CI16) inoculated simultaneously with the target strain (*B. cereus* UW85 GFP). (B) Variable strains inoculated and grown for 6 hours prior to inoculating target strains. Growth equal to the no-variable control would have a value of 1 (shown as the black dotted line). Error bars denote the standard deviation from 2 replicates.

### Community combination screening of a 10-member rhizosphere community on the μCI device

To test the ability to measure effects of various bacterial species on the target strain, we selected ten strains from the THOR rhizosphere bacteria collection (45) and *B. cereus* UW85 GFP as the target strain. Ten strains result in 120 possible three-way combinations, and thus, full combinatorial space coverage was achieved using two μCI devices (88-combination wells/device). Geometrical constraints led to more than one repetition of some combinations in each experiment. Overnight liquid cultures of variable strains were inoculated at OD 0.01 into the variable wells and cultured for 6 h followed by addition of the target strain (*B. cereus* UW85 GFP) at OD 0.01 in the combination wells. A control device containing only buffer (PBS) in the variable wells was included in each experiment for growth normalization (Fig. S5). The devices were then incubated at RT for 2 days with fluorescence images captured on both day 1 and day 2. Large differences in growth across strain combinations were evident one day after inoculation (Fig. 5A-C, Table S1) with both increased and decreased growth relative to the no-variable strain control (Fig. 5C). We observed similar target strain growth trends across different strain combinations for day 1 and day 2 (R^2^ = 0.8584), except the fluorescence intensity of the target strain was generally brighter on day 1 across the board (Fig. 5D), suggesting that GFP expression diminished as culture media saturation was achieved.

**Fig. 5.**
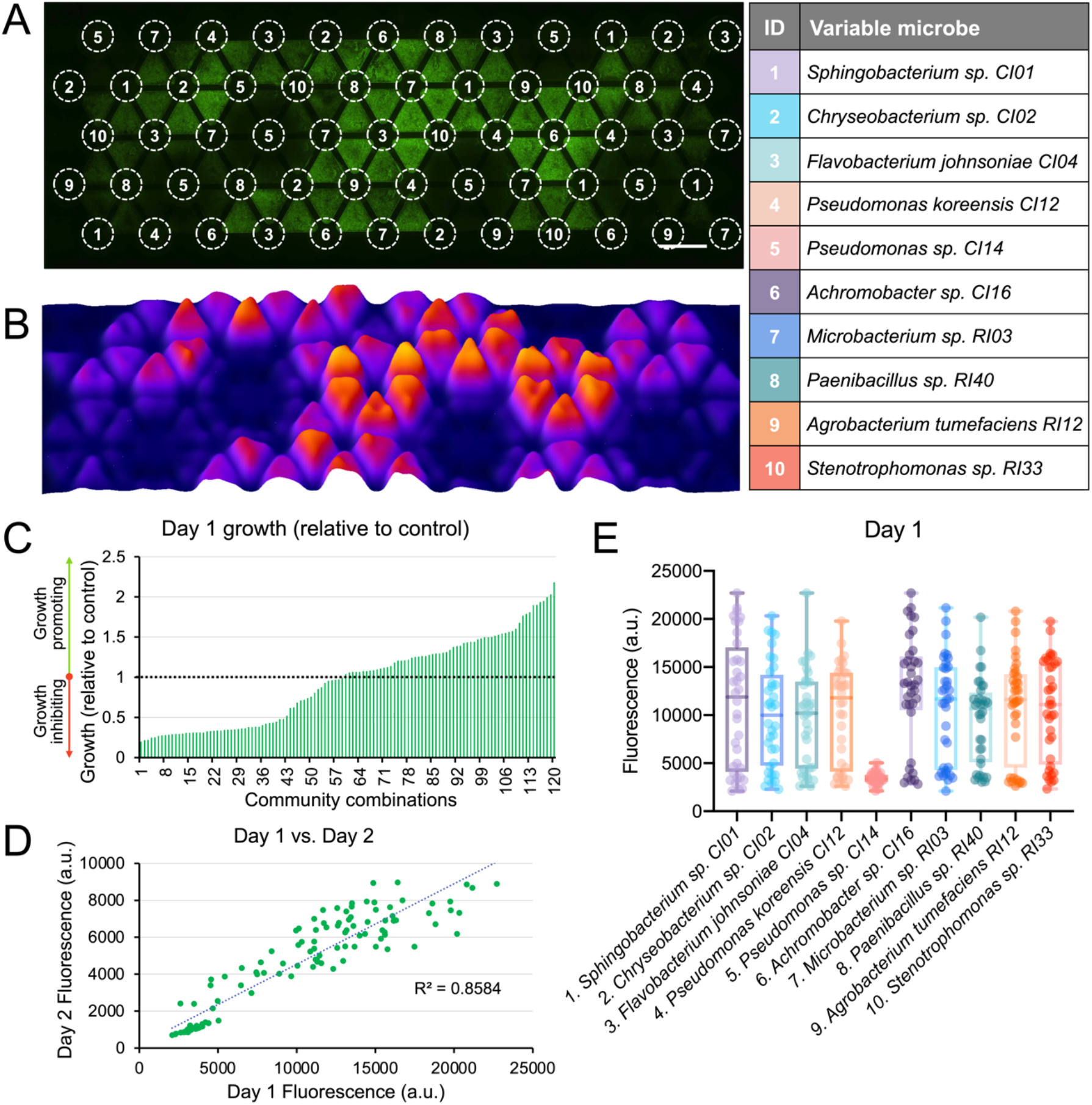
Community combination screening of a 10-member rhizosphere community on the μCI device. (A) Fluorescence microscopy images of target strain *B. cereus* UW85 GFP on the μCI device after co-culture with a 10-member rhizosphere community. (B) Surface “landscape” plot of (A) with fluorescence intensity represented in the *z*-axis. (C) Quantified growth (relative to no-variable control) of the target strain when co-cultured with each microbial community. Data is organized in increasing order. Growth equal to the no-variable control would have a value of 1 (shown as the black dotted line). (D) Linear regression of target strain growth on day 1 vs. day 2 across all 3-member variable communities. (E) Target strain growth sorted by variable microbial communities which contain the indicated variable strain (n=3).

To gain insight into the growth promoting/inhibiting trends of each strain within the community, we sorted the growth data by communities which contain the indicated variable strain (Fig. 5E). While each data point represents the integrated effect of a three-member variable community on the target strain, growth promoting/inhibiting trends for individual variable strains were observed. For instance, presence of *Pseudomonas* sp. CI14 (microbe 5) generally reduced growth across all strain combinations, whereas strain combinations containing *Achromobacter sp. CI16* (microbe 6) often enhances growth of *B. cereus* UW85 (Fig. 5E). Variable strains *Pseudomonas* sp. CI14, *Achromobacter* sp. CI16, and *Paenibacillus* sp. RI40 resulted in a narrower growth distribution of the target strain suggesting that they modulated growth more strongly than strains that exhibit broader target strain growth distributions (Fig. 5E). Time-lapse microscopy of the target strain in the community over time revealed similar strain-specific growth-modulating trends (Fig. S6), with *Pseudomonas* sp. CI14 showing the greatest growth inhibition.

Interestingly, autofluorescence images in the UV channel (excitation 390 nm, emission 440 nm) showed that *Pseudomonas* sp. CI14 secretes a compound that fluoresces blue in this channel (Fig. S7), which we suspect may potentially have antimicrobial activity. To test whether the secretion of the fluorescent blue compound is modulated by co-culture, we cultured *Pseudomonas* sp. CI14 alone or in co-culture with *B. cereus* UW85 GFP, *Chryseobacterium* sp. CI02, *Flavobacterium johnsoniae* CI04, and *Achromobacter* sp. CI16. However, we found that the blue autofluorescence was not affected by co-culture with any of the strains (Fig. S8), indicating that its secretion was not influenced by interactions with these strains. Interestingly, certain microbe combinations consist of strains that are not strongly inhibitory towards *B. cereus* UW85 alone but show strong inhibitory activity as a 3-member combination (combinations 2,3,4 and 1,7,9 for instance) (Table S1), demonstrating the capability of the μCI device in identifying potential emergent microbial community phenotypes. To summarize, we successfully performed a combinatorial screen of all 120 possible 3-member combinations of 10 microbes from a rhizosphere community on the μCI device, and identified a strain which played a strong inhibitory role towards *B. cereus* UW85 (*Pseudomonas* sp. CI14), as well as 3-member communities which display potential emergent phenotypes.

## Discussion

Here we describe the μCI platform for systematic study of microbial community interactions. The μCI device provides measurements of chemical and microbial interactions within three-member subsets of a community, and was designed to be easy to operate with only pipettes and a fluorescent microscope, making it accessible to non-engineering labs. The μCI device demonstrated its utility by rapidly identifying a rhizosphere strain, *Pseudomonas* sp. CI14, as highly inhibitory to other community members. The same conclusion was reached previously with laborious, pairwise experiments on agar plates and a hierarchy model (45), whereas the present experiment required only ∼2 hours for setup and 2 days for co-culture and readout. The μCI device also makes it easy for discovering positive interactions such as growth stimulators. These are underrepresented interactions in the literature since there is a lack of simple-to-use, high-throughput platforms for identifying these interactions, so their prevalence and significance in nature is relatively underexplored. Also, as evidenced in our rhizosphere microbe co-culture data, the μCI device enables the identification of emergent properties between members of the same community.

A current limitation of the μCI device is the requirement for a fluorescently labeled target strain, which reduces its utility in studying microbes for whom a fluorescent protein reporter is not available. This also limits the platform’s ability to assay microbial communities that require anaerobic conditions (such as gut communities), as most traditional protein fluorophore reporters do not function in anaerobic conditions (51) It will be of interest to develop alternative microbial growth quantification strategies such as chemical reporter dyes or non-fluorescent imaging and image processing methods to assay microbes without a fluorescent reporter. We also envision that the μCI device will be a good platform for screening combinations of candidate microorganisms for probiotics for treatment of diseases that are influenced or caused by microorganisms, or the study of the resistance or susceptibility of communities to invaders.

Perhaps the greatest power of the μCI device lies in its potential for identifying regulatory networks in communities. By replacing the constitutive GFP construct with one containing a promoter of interest, hundreds of chemicals or organisms could be screened simultaneously to discover regulatory factors. Since genes that are uniquely important for survival in a community are largely undiscovered or poorly characterized, this capability could revolutionize the process of assigning functions for community-specific genes and identifying regulatory cascades. In addition, gene clusters responsible for synthesis of small molecules are often silent. That capacity for rapid screening of potential inducers of expression could facilitate discovery of new bioactive molecules with potential for applications in agriculture and medicine.

## Materials and Methods

### Device design and fabrication

Design of the μCI devices was done using computer-aided design (CAD) software (AutoCAD, Autodesk Inc), and fabricated using a computer numerical control (CNC) milling machine (Tormach PCNC 770) with polystyrene as the device material as previously described (52) The μCI device design employs a two-layer fabrication approach. Namely, the variable wells and combination wells are respectively milled out of 1.2-mm and 2-mm polystyrene, then bonded together via acetonitrile-assisted heat bonding. In this configuration the variable wells are on one side of the device and the combination wells on the other side, facing opposite directions. Holders for the device (to keep the device suspended) was fabricated with polylactic acid (PLA) polymer using a fused deposition modeling (FDM) 3D printer (Ultimaker 3) and placed in an OmniTray single-well plate (Thermo Scientific Nunc). Luria-Bertani (LB) agar was prepared and sterilized in an autoclave, then pipetted into the μCI device using a 20-μL pipet while warm (15 μL of LB agar was pipetted into the variable wells first and allowed to solidify, followed by flipping the device over and pipetting 17 μL of LB agar into the combination wells.)

### Microbial Culture and Antibiotic Testing

The 10 rhizosphere bacterial strains and *B. cereus* UW85 GFP fluorescent reporter strain were cultured as reported previously (45). Bacteria were grown in liquid culture overnight in 50% strength tryptic soy broth (TSB) at 28°C. Turbidity measurements of the liquid cultures were performed using an ELISA reader at 600 nm. Aliquots of cultures of each strain were centrifuged at 6,000 × g for 6 min, the supernatant was discarded, and the bacteria pellet was resuspended in PBS to an equivalent OD 600 of 0.01. For the variable wells of the μCI device, 3 μL of antibiotics in PBS (for antibiotic testing experiments) or 0.01 OD bacteria suspension (variable strains for microbial community experiments) was dispensed on the agar surface using a 10 μL pipet. The device was then flipped over, allowing the combination wells to face upwards. Three microliters of 0.01 OD bacteria suspension (the target strains) were dispensed on the agar surface of the combination wells using a 10 μL pipet (either 6 h after or immediately after inoculating the variable strains). Care was taken to not poke holes on the agar surface with the pipet tip (which can affect image analysis). The devices were then placed in a laminar flow hood for 15 min to allow the liquid on the agar to air dry. The devices were placed on a holder in an OmniTray, which was then sealed on the edges using parafilm to prevent evaporation and drying of the agar. Devices were incubated at RT for 2 days and bacterial growth was monitored using an epifluorescence microscope at set time points.

### Fluorescence Microscopy

Imaging of microbial growth was performed using a Nikon Ti-Eclipse inverted epifluorescence microscope equipped with a motorized *xy* stage. A stitched montage image of the whole device was acquired using a 2× objective in bright-field and two fluorescence excitation channels: 390 nm (blue), and 485 nm (green).

### Layout Generation and Image Processing

A custom layout generator script was written in MATLAB (The MathWorks, Natick, MA) to generate the layout for plating microbes in the variable wells. The script attempts to maximize combinatorial coverage on the device and minimize repeats. This was achieved by using a random number generator to determine which microbe would reside in each variable well. The number of combination wells containing each microbe was recorded and used to determine the probability distribution for placing each microbe in future wells. Microbes were then plated according to the generated layout. The custom MATLAB script is freely available at https://github.com/mmblab/CommunityFitnessLandscape. Following culture and image acquisition, the images were then loaded into Fiji (https://fiji.sc/). A custom ROI array was created using the ROI manager of Fiji to measure the mean fluorescence intensity of each combination well and the three edges of the combination well. The measured fluorescence intensity values were then exported to a CSV file, then imported into MATLAB. We wrote a custom MATLAB script to map each ROI datapoint to the associated strain combination indicated by the custom layout generated previously.

### Diffusion Experiments

Six-well μCI devices with various dimensions ranging from a pitch of 4.5, 5.5, 6, and 6.5 mm (between 2 variable wells) were fabricated and filled with agar as described above. The volume of agar in each well was scaled proportionally according to the volume of the device. A 10 μg/mL solution of Rhodamine 6G (Millipore Sigma) solution in PBS was pipetted onto the surface of the agar in the variable well. Diffusion was monitored at 30-min intervals for 12 hours via time-lapse microscopy using a Nikon Ti-E inverted epifluorescence microscope with the 560 nm excitation channel. A linear ROI was drawn in Fiji starting from the inner edge of the triangular combination well and ending at the midpoint of the outer side of the well to quantify the diffusion profile as reflected in the fluorescence intensity.

### Finite Element Modeling

Modeling of diffusion in the μCI device was performed using COMSOL Multiphysics software (COMSOL, Burlington, MA). A 3-dimensional model of the agar within the μCI device was drawn using AutoCAD and imported into COMSOL. For diffusion analysis, the *Transport of Diluted Species in Porous Media* module was used. The diffusion source was set at the top surface of the agar in the variable well, with a diffusion coefficient of 1 × 10^−6^ cm^2^ ⋅s^−1^ (similar to the diffusion coefficient of antibiotics in agar as previously reported (53)). The porosity of the agar was set to 0.9805 (the porosity of 2% agarose gel as previously reported (54). A time-dependent simulation was performed with 30-min intervals across 12 hours corresponding to the Rhodamine 6G diffusion experiment.

## Acknowledgments

We would like to thank Dr. Jay Warrick, and Dr. Scott Berry for helpful input and suggestions for the project. This study was funded by the National Institutes of Health (NIH P30CA014520, R35 GM124774, R01 EB030340), the U.S. Army Research Laboratory and the U.S. Army Research Office under contract/grant W911NF1910269, the Office of the Vice Chancellor for Research and Graduate Education SEED program, and the State of Wisconsin.

## Author Contributions

D.S.J., W.E.W., and G.L.L. designed and performed experiments. L.J.B., J.Y., A.H., M.F.G.D., and O.S.V. provided insight into the study and assisted in experimental design. D.J.B., J.H., and G.L.L. oversaw the study, edited the manuscript, and obtained funding for the project. D.S.J. and W.E.W. wrote the manuscript with input from all authors.

