## Supporting Information for "Microbial Community Interactions on a Chip"

**Supporting Information for**
### Microbial Community Interactions on a Chip

Duane. S. Juang[a], Wren E. Wightman[a], Gabriel L. Lozano[b,g], Layla J. Barkal[a], Jiaquan Yu[a], Manuel F. Garavito[b], Amanda Hurley[b], Ophelia S. Venturelli[c,d,e], Jo Handelsman[b], and David J. Beebe[a,f]

[a]Department of Biomedical Engineering, University of Wisconsin-Madison, Madison, WI, USA
[b]Wisconsin Institute for Discovery and Department of Plant Pathology, University of Wisconsin-Madison, Madison, WI, USA
[c]Department of Biochemistry, University of Wisconsin-Madison, Madison, WI, USA
[d]Department of Bacteriology, University of Wisconsin-Madison, Madison, WI, USA
[e]Department of Chemical & Biological Engineering, University of Wisconsin-Madison, Madison, WI, USA
[f]Department of Pathology and Laboratory Medicine, University of Wisconsin-Madison, Madison, WI, USA
[g]Current address: Division of Infectious Disease, Department of Pediatrics, Boston Children's Hospital, Harvard Medical School, Boston, MA, USA

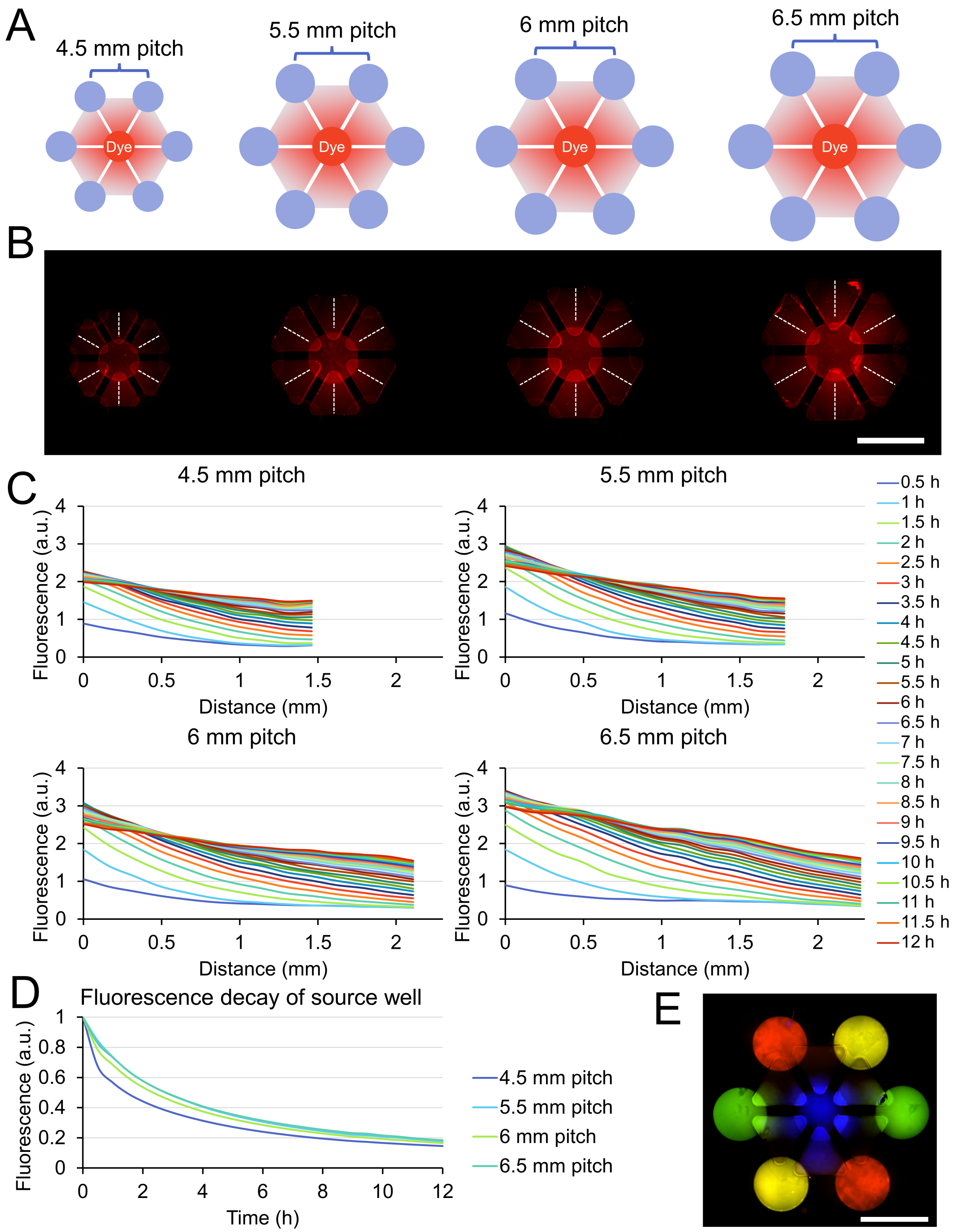


**Fig. S1.** Soluble factor diffusion in the µCI device. (A) Diffusion was measured by adding a fluorescent small molecule dye (Rhodamine 6G) to the center variable well and allowing for passive diffusion over a period of 12 hours. (B) Fluorescence images of Rhodamine 6G diffusion from the center variable well. Scale bar: 5 mm. (C) Mean fluorescence diffusion profiles across the combination wells (dotted white line in (B)) over a 12-hour period. (D) Fluorescence decay of the center variable (source) well over a 12-hour period. (E) 4-color combinations were accomplished using 4 cell tracking dyes (Blue: cell tracker blue, green: cell tracker green, red: cell tracker red, yellow: cell tracker deep red). Scale bar: 5 mm.

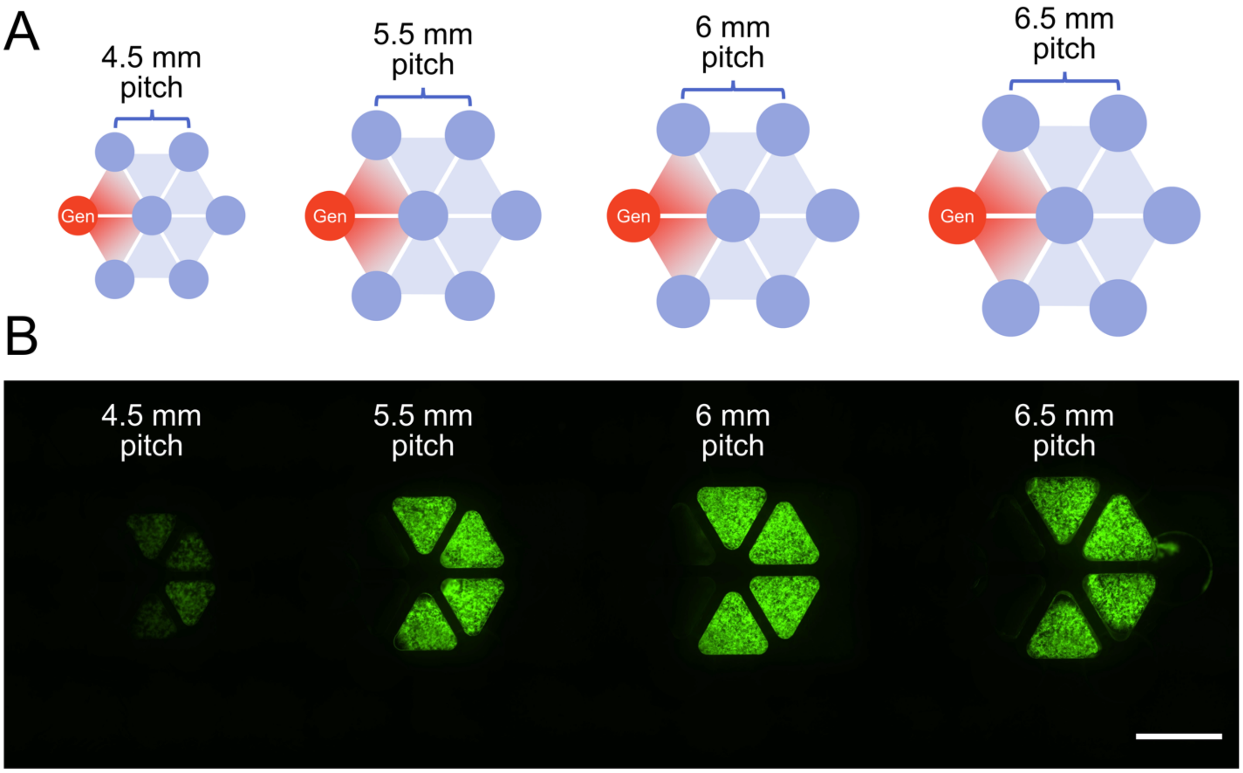


**Fig. S2.** Antibiotic diffusion in the µCI device across different well pitches. (A) Experimental layout of gentamicin diffusion experiment. Gentamicin was added at a concentration of 80 µg/mL into the left-most variable well. The target strain (*B. cereus* UW85 GFP) was inoculated into the combination wells and cultured for 1 day. (B) Fluorescence microscopy images of the µCI device with 4 different well pitches. Scale bar: 5 mm.

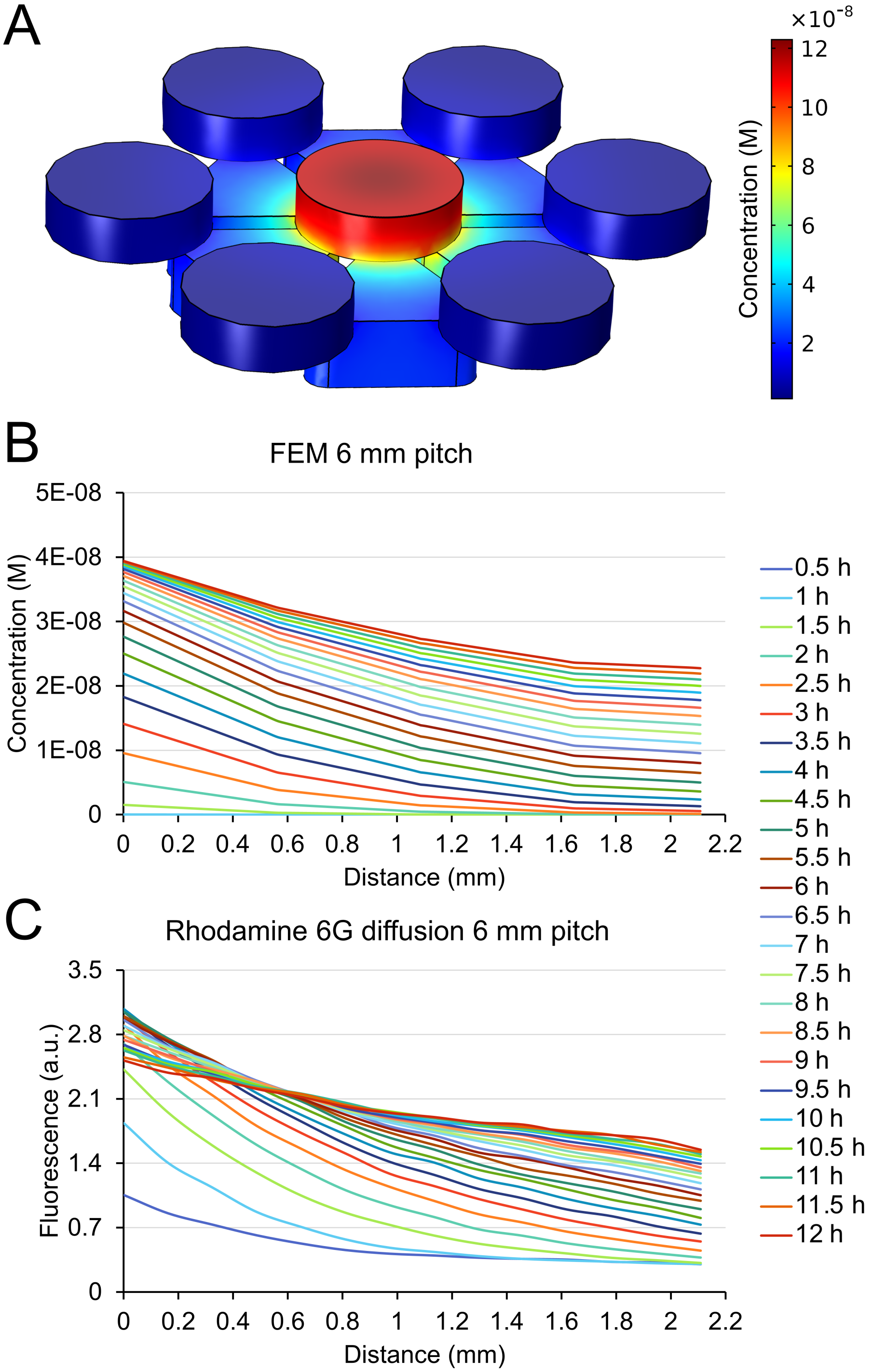


**Fig. S3.** Finite element modeling (FEM) of small molecule diffusion in the 6 mm pitch µCI device. (A) 3D COMSOL model of diffusion within the µCI device. (B) FEM diffusion profile across the agar surface of the combination well (where the target microbes reside). (C) Fluorescence diffusion profile of rhodamine 6G across the combination well (same as Fig. S1C, panel 3).

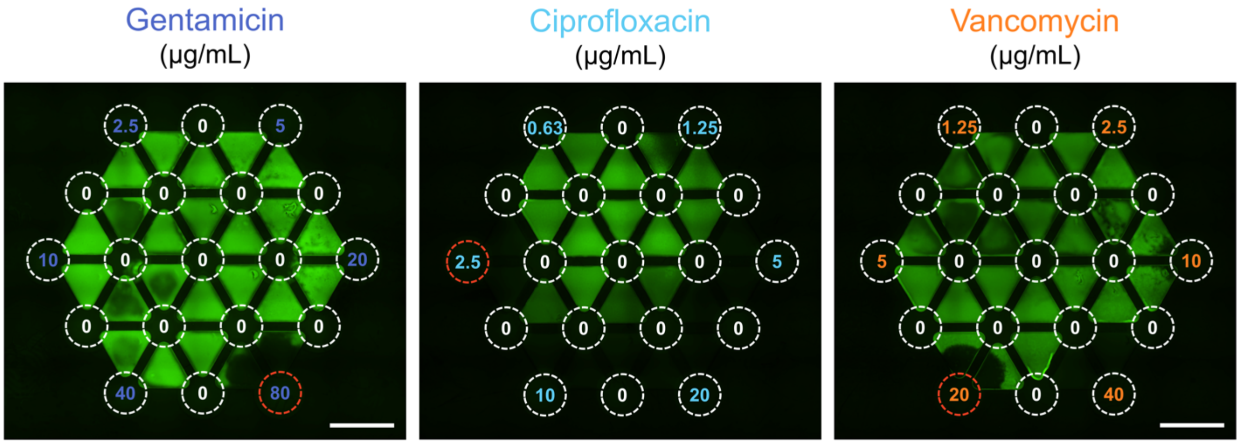


**Fig. S4.** Antibiotic susceptibility testing of three antibiotics (gentamicin, ciprofloxacin, and vancomycin) at various concentrations to determine their minimum inhibitory concentration (MIC) on the µCI device. MIC conditions circled in red. Scale bar: 5 mm.

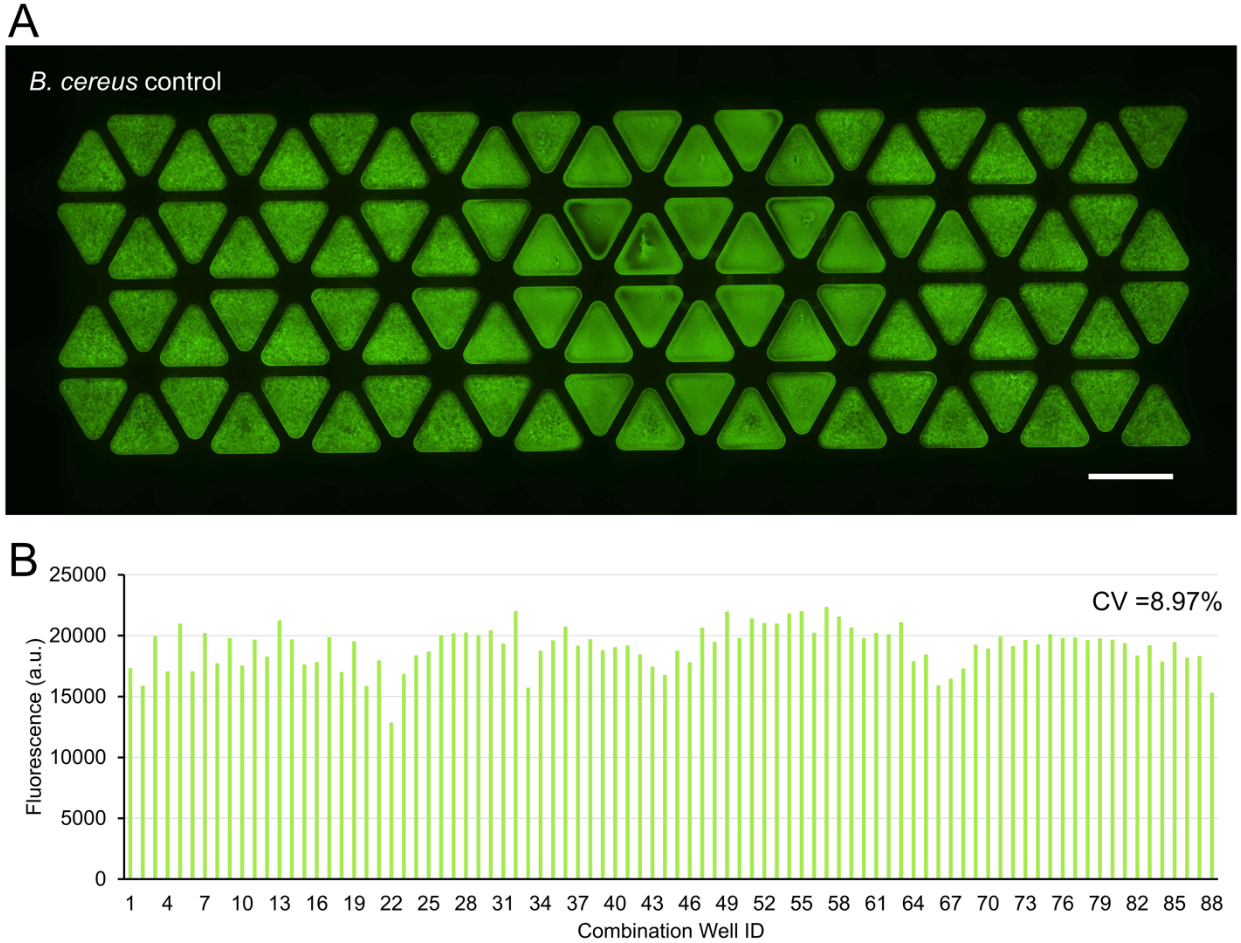


**Fig. S5.** Growth of target strain (*B. cereus* UW85 GFP) without the presence of variable strains. (A) Fluorescence image, and (B) quantified fluorescence of each well of the µCI device inoculated with the target strain in the combination wells. Scale bar: 5 mm.

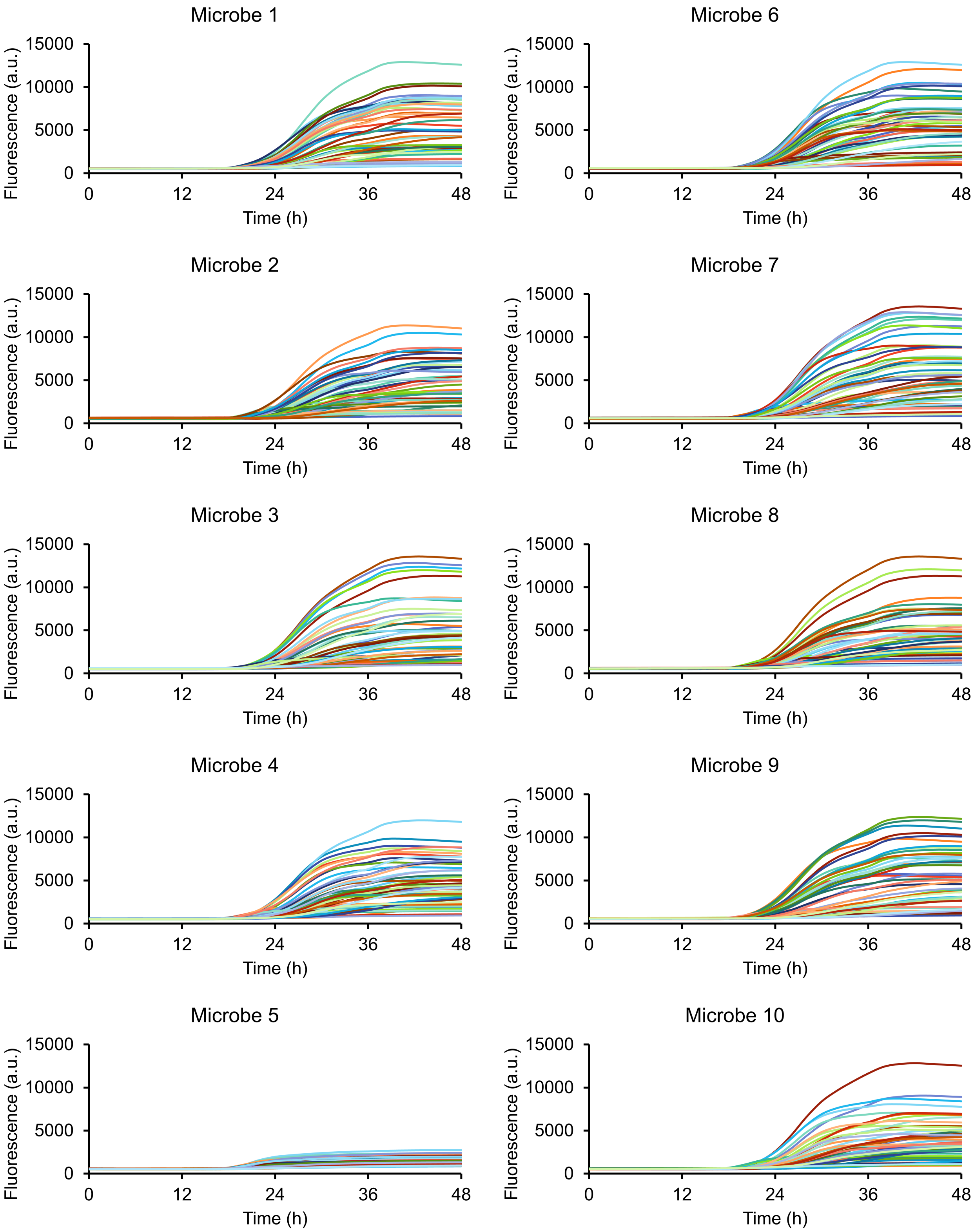


**Fig. S6.** 48-hour time-lapse growth curves of target strain sorted by variable microbial communities which contain the indicated variable strain.

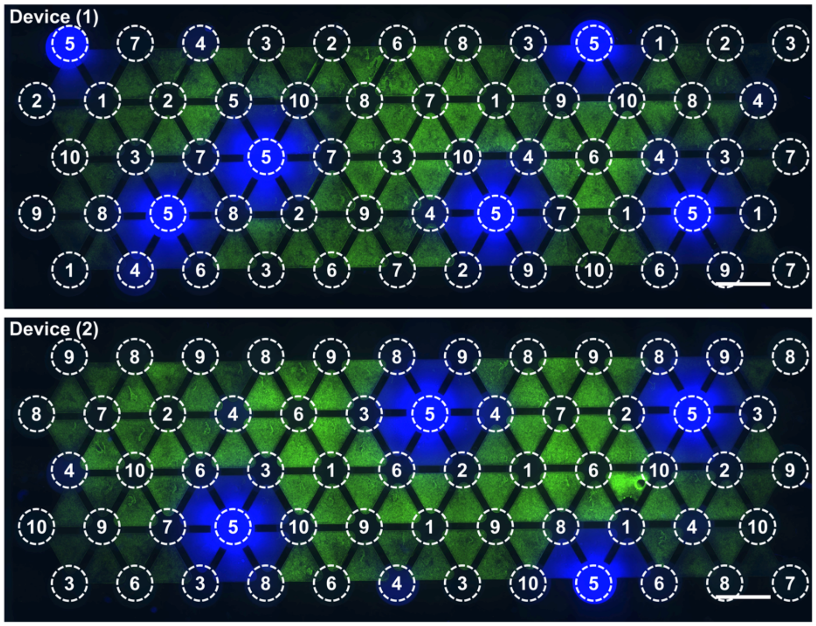


| ID | Variable microbe |
|---|---|
| 1 | *Sphingobacterium sp. CI01* |
| 2 | *Chryseobacterium sp. CI02* |
| 3 | *Flavobacterium johnsoniae CI04* |
| 4 | *Pseudomonas koreensis CI12* |
| 5 | *Pseudomonas sp. CI14* |
| 6 | *Achromobacter sp. CI16* |
| 7 | *Microbacterium sp. RI03* |
| 8 | *Paenibacillus sp. RI40* |
| 9 | *Agrobacterium tumefaciens RI12* |
| 10 | *Stenotrophomonas sp. RI33* |

**Fig. S7.** Fluorescence microscopy image of community combination screening of a 10-member rhizosphere community on the µCI device. Green: target strain (*B. cereus* UW85 GFP), Blue: autofluorescence. Scale bar: 5 mm.

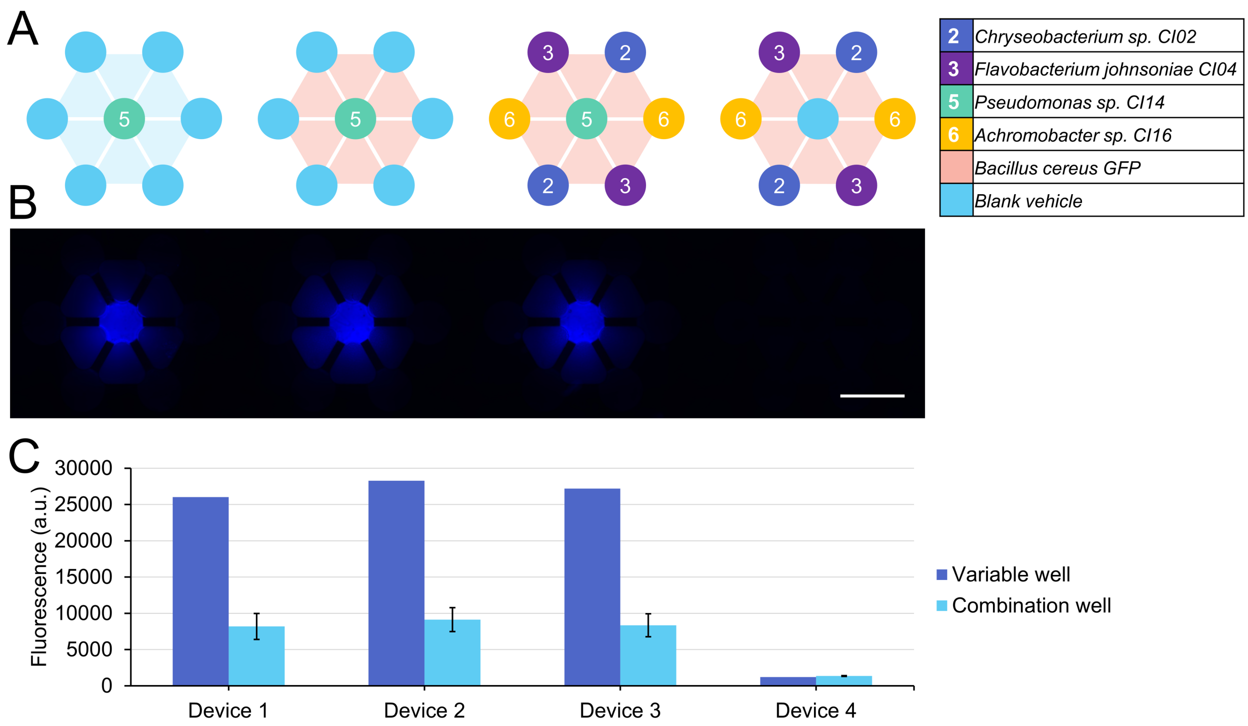


**Fig. S8.** Secretion of fluorescent blue compound by *Pseudomonas* sp. CI14 in monoculture or co-culture. (A) Experimental layout. (B) Fluorescence image of fluorescent blue compound secreted by *Pseudomonas* sp. CI14. (C) Fluorescence of the center variable well and surrounding combination wells quantified from (B).

**Table S1.** Growth of *B. cereus* UW85 GFP target strain in co-culture with three-member variable microbial communities (data from Fig. 5C).

| ID | Variable microbe |
|---|---|
| 1 | *Sphingobacterium* sp. CI01 |
| 2 | *Chryseobacterium* sp. CI02 |
| 3 | *Flavobacterium johnsoniae* CI04 |
| 4 | *Pseudomonas koreensis* CI12 |
| 5 | *Pseudomonas* sp. CI14 |
| 6 | *Achromobacter* sp. CI16 |
| 7 | *Microbacterium* sp. RI03 |
| 8 | *Paenibacillus* sp. RI40 |
| 9 | *Agrobacterium tumefaciens* RI12 |
| 10 | *Stenotrophomonas* sp. RI33 |

| Variable community combination | Target strain growth (relative to control) |
|---|---|
| 1,5,7 | 20.02% |
| 1,2,5 | 21.84% |
| 1,5,10 | 22.23% |
| 2,3,4 | 24.99% |
| 3,5,9 | 25.22% |
| 4,5,6 | 27.13% |
| 2,3,5 | 27.74% |
| 1,5,9 | 28.07% |
| 5,6,9 | 28.60% |
| 4,5,8 | 29.09% |
| 5,9,10 | 29.36% |
| 5,8,9 | 29.49% |
| 2,5,9 | 29.71% |
| 5,6,10 | 30.83% |

| | |
|---|---|
| 1,3,5 | 30.92% |
| 1,5,6 | 31.14% |
| 5,7,9 | 31.32% |
| 5,8,10 | 31.39% |
| 2,4,5 | 31.41% |
| 3,4,5 | 32.56% |
| 4,5,9 | 33.14% |
| 1,7,9 | 33.31% |
| 5,6,8 | 33.66% |
| 1,5,8 | 33.66% |
| 3,5,8 | 34.28% |
| 4,5,10 | 34.88% |
| 4,5,7 | 34.91% |
| 3,5,7 | 35.31% |
| 5,7,8 | 35.66% |
| 2,5,10 | 36.37% |
| 2,5,8 | 36.39% |
| 5,6,7 | 37.91% |
| 1,4,5 | 38.09% |
| 3,5,10 | 38.37% |
| 2,5,7 | 38.44% |
| 5,7,10 | 40.32% |
| 3,5,6 | 42.15% |
| 1,3,7 | 43.43% |
| 3,4,7 | 43.71% |
| 1,2,10 | 44.79% |
| 1,4,8 | 47.75% |
| 2,5,6 | 48.18% |
| 3,8,10 | 52.13% |
| 1,2,8 | 61.85% |

| | |
|---|---|
| 2,4,8 | 62.38% |
| 1,2,7 | 68.29% |
| 7,8,10 | 71.31% |
| 3,4,8 | 71.83% |
| 2,9,10 | 74.13% |
| 1,2,3 | 75.89% |
| 2,3,10 | 80.74% |
| 2,4,7 | 85.25% |
| 2,3,9 | 87.66% |
| 1,8,10 | 92.65% |
| 3,9,10 | 95.55% |
| 4,8,10 | 96.47% |
| 1,3,8 | 97.01% |
| 4,9,10 | 97.24% |
| 3,6,8 | 99.09% |
| 2,3,7 | 104.21% |
| 3,8,9 | 105.92% |
| 2,8,10 | 106.60% |
| 3,6,10 | 106.67% |
| 2,6,8 | 106.68% |
| 3,6,7 | 107.53% |
| 4,7,9 | 107.72% |
| 2,3,8 | 108.35% |
| 4,8,9 | 108.63% |
| 1,8,9 | 109.95% |
| 8,9,10 | 110.98% |
| 7,8,9 | 112.05% |
| 6,7,8 | 112.50% |
| 2,3,6 | 114.11% |
| 1,4,6 | 118.31% |

| | |
|---|---|
| 2,8,9 | 120.62% |
| 6,7,9 | 120.65% |
| 7,9,10 | 121.04% |
| 1,7,10 | 121.80% |
| 2,4,10 | 124.33% |
| 4,7,8 | 125.50% |
| 4,6,8 | 125.57% |
| 3,4,9 | 126.30% |
| 2,4,6 | 126.38% |
| 2,6,9 | 127.74% |
| 3,6,9 | 129.02% |
| 1,7,8 | 129.53% |
| 3,7,8 | 129.76% |
| 1,3,9 | 130.57% |
| 4,6,10 | 131.21% |
| 1,4,9 | 134.37% |
| 4,6,9 | 137.75% |
| 1,3,4 | 138.73% |
| 2,4,9 | 139.27% |
| 3,7,9 | 142.65% |
| 4,7,10 | 143.45% |
| 6,8,10 | 144.24% |
| 2,7,8 | 144.27% |
| 6,9,10 | 147.45% |
| 4,6,7 | 148.30% |
| 1,6,10 | 149.67% |
| 1,3,10 | 149.76% |
| 3,4,6 | 150.07% |
| 1,4,10 | 151.45% |
| 2,7,10 | 152.94% |

| | |
|---|---|
| 2,7,9 | 154.49% |
| 3,4,10 | 155.15% |
| 6,7,10 | 156.51% |
| 3,7,10 | 157.69% |
| 6,8,9 | 160.52% |
| 1,2,4 | 167.82% |
| 2,6,7 | 176.60% |
| 1,2,9 | 178.84% |
| 2,6,10 | 180.73% |
| 1,9,10 | 189.60% |
| 1,4,7 | 190.05% |
| 1,6,8 | 193.79% |
| 1,2,6 | 195.19% |
| 1,6,9 | 199.81% |
| 1,6,7 | 203.26% |
| 1,3,6 | 218.12% |
